# An Empirical Demonstration of Unsupervised Machine Learning in Species Delimitation

**DOI:** 10.1101/429662

**Authors:** Shahan Derkarabetian, Stephanie Castillo, Peter K. Koo, Sergey Ovchinnikov, Marshal Hedin

**Author notes:** Corresponding author: Shahan Derkarabetian Department of Organismic and Evolutionary Biology Museum of Comparative Zoology Harvard University Cambridge, MA 02138.

## Abstract

One major challenge to delimiting species with genetic data is successfully differentiating species divergences from population structure, with some current methods biased towards overestimating species numbers. Many fields of science are now utilizing machine learning (ML) approaches, and in systematics and evolutionary biology, supervised ML algorithms have recently been incorporated to infer species boundaries. However, these methods require the creation of training data with associated labels. Unsupervised ML, on the other hand, uses the inherent structure in data and hence does not require any user-specified training labels, thus providing a more objective approach to species delimitation. In the context of integrative taxonomy, we demonstrate the utility of three unsupervised ML approaches, specifically random forests, variational autoencoders, and t-distributed stochastic neighbor embedding, for species delimitation utilizing a short-range endemic harvestman taxon (Laniatores, *Metanonychus*). First, we combine mitochondrial data with examination of male genitalic morphology to identify a priori species hypotheses. Then we use single nucleotide polymorphism data derived from sequence capture of ultraconserved elements (UCEs) to test the efficacy of unsupervised ML algorithms in successfully identifying a priori species, comparing results to commonly used genetic approaches. Finally, we use two validation methods to assess a priori species hypotheses using UCE data. We find that unsupervised ML approaches successfully cluster samples according to species level divergences and not to high levels of population structure, while standard model-based validation methods over-split species, in some instances suggesting that all sampled individuals are distinct species. Moreover, unsupervised ML approaches offer the benefits of better data visualization in two-dimensional space and the ability to accommodate various data types. We argue that ML methods may be better suited for species delimitation relative to currently used model-based validation methods, and that species delimitation in a truly integrative framework provides more robust final species hypotheses relative to separating delimitation into distinct “discovery” and “validation” phases. Unsupervised ML is a powerful analytical approach that can be incorporated into many aspects of systematic biology, including species delimitation. Based on results of our empirical dataset, we make several taxonomic changes including description of a new species.

---

Modern species delimitation is becoming increasingly objective relying on, for example, statistical thresholds and/or clustering algorithms to identify species in multivariate morphological space (e.g., Ezard et al. 2010; Seifert et al. 2014), or using the multispecies coalescent to identify the boundary between population and species level divergences using genetic data (e.g., Yang and Rannala 2010). Similarly, species delimitation is becoming increasingly integrative, combining multiple data types in a reciprocally-illuminating framework providing more robust final species hypotheses (Dayrat 2005; Schlick-Steiner et al. 2010). The empirical process of delimiting species has been portrayed by some authors as occurring in two separate phases (Carstens et al. 2013): a discovery phase where a priori hypotheses are formed based on one or more data types, followed by a validation phase where species hypotheses are further tested using an independent dataset, typically nuclear genetic data. Of utmost interest in using genetic data in species delimitation, whether as validation or otherwise, is successfully distinguishing population structure from species level divergences. Recently, Sukumaran and Knowles (2017) demonstrated that the multispecies coalescent model will support population level divergences, an assertation previously demonstrated empirically (e.g., Niemiller et al. 2012; Hedin et al. 2015).

Across many fields of science, a great deal of attention has been given to machine learning (ML) approaches, where an algorithm can be trained to make future decisions without user input. Recently, ML methods like random forest (RF; Breiman 2001) have been incorporated into systematics and evolutionary biology, with applications in barcoding (e.g., Austerlitz et al. 2010), environmental DNA metabarcoding (e.g., Cordier et al. 2018), population genetics (e.g., Schrider and Kern 2016; Schrider and Kern 2018), and predicting cryptic diversity (Espíndola et al. 2016). Most relevant here is the use of RF in phylogeographic model selection (Pudlo et al. 2016; Smith et al. 2017) and speciation/species delimitation (Pei et al. 2018; Smith and Carstens 2018) where it can be used as a validation tool distinguishing among multiple user-specified models given a priori information about the training data. Similarly, non-RF ML approaches have been used to model biogeographic processes (Sukumaran et al. 2015). In these examples a *supervised* ML approach is used, where simulated datasets based on user-specified priors are used as training data, and a classifier is built to choose among different models or species hypotheses given observed data. For example, the recently developed RF-based species delimitation program CLADES (Pei et al. 2018) approaches species delimitation as a classification issue. Here, a two-species model with varying divergence times and population sizes, with or without migration, is used to simulate the training datasets for classifier construction. Multiple population genetic summary statistics are computed for labeled training data and observed data with species hypotheses defined a priori. These statistics are used as variables to determine support for a priori species distinctiveness in the observed data.

While supervised approaches are indeed powerful, unsupervised ML may also be a useful approach to aid in species delimitation using the inherent structure in the data to cluster samples. Unsupervised ML can be conducted without a priori hypotheses regarding the underlying evolutionary model, population parameters, number of species, species assignment, or levels of parameter divergence needed to classify samples as different species. In unsupervised RF, the training data is a synthetic dataset based on the observed data representing the null hypothesis of no structure, and a classifier is built to distinguish the synthetic and observed datasets, thus uncovering underlying structure (if present) in the observed data. Many unsupervised ML algorithms for high-dimensionality data intrinsically perform reduction to a lower dimensional space, where the underlying data structure can be visualized. For example, Oltaenu et al. (2013) take an unsupervised ML approach to visualizing and clustering barcode data via nonlinear dimension reduction and projection methods using multidimensional scaling and self-organizing maps. They show that these approaches successfully clustered named and unnamed species and suggested the possibility of undescribed species.

Many ML algorithms can be executed in an unsupervised manner, and while dimensionality reduction methods like principal components analyses and clustering algorithms like k-means are widely considered to be ML, we focus on three unsupervised ML approaches chosen to represent a diversity of ML algorithm types including one that has yet to be used in the field of systematics (Table 1): Random Forests (RF; Breiman 2001), Variational Autoencoders (VAE; Kingma and Welling 2013), and t-Distributed Stochastic Neighbor Embedding (t-SNE; van der Maaten and Hinton 2008). RF is an ensemble learning method that relies on classification trees and tree bagging (Breiman 1996; 2001). In RF most importantly, in a given classification tree if two samples appear at the same terminal node their “proximity score” is increased by one. Proximity scores for all pairs are averaged over bootstrap replicates to produce a final proximity matrix, which can be used in multidimensional scaling (MDS) and clustering. A Variational Autoencoder is a Bayesian approach that learns a distribution of the data using latent variables. It does so in two stages: 1) inference of the posterior distribution of latent variables and 2) generation of data sampled from a given set of latent values. Both stages are approximated by neural networks and optimized simultaneously via unsupervised learning. Widely-used in diverse fields (e.g., Bauer et al. 2015; Yoshida et al. 2016; Mallet et al. 2017), t-SNE is a non-linear dimensionality reduction algorithm that attempts to preserve probability distributions of distances among samples within a cluster but repels samples that are in different clusters in lower-dimensional space.

**Table 1.** Comparison of unsupervised machine learning methods used in this study.

| Method | Purpose | Approach used | General algorithm | Relevant output |
| --- | --- | --- | --- | --- |
| Random Forest (RF) | Classification and regression | Supervised<br>Unsupervised | Ensemble method that grows many classification trees based on training data, runs input data down trees, and the classification with the most votes is chosen. | Proximity matrix |
| Variational Autoencoder (VAE) | Generative model | Unsupervised | Compresses data through multiple encoding layers into latent variables, then un-compresses latent variables through multiple decoder layers into reconstructed data. Learns the marginal likelihood distribution of the data using latent variables. | Latent variables (two-dimensional encoding) |
| t-Distributed Stochastic Neighbor Embedding (t-SNE) | Data embedding and visualization | Unsupervised | Constructs probability distribution of sample pairs, then minimizes divergence between high dimensional space and low dimension embedding, such that similar pairs are embedded nearby while dissimilar pairs are repelled. | Low dimensional embedding |

The purpose of this study was, in the context of integrative taxonomy, to explore and demonstrate the utility of unsupervised ML approaches in aiding species delimitation through successful identification of clusters corresponding to species, as corroborated by other traditional methods. First, in the discovery phase, we combine phylogenetic analysis of mitochondrial cytochrome oxidase subunit I (COI) and examination of morphology to generate a priori species hypotheses. Then, using single nucleotide polymorphisms (SNPs) derived from sequence capture of ultraconserved elements (UCEs) we demonstrate the ability of unsupervised ML approaches to successfully cluster identified a priori species, comparing three unsupervised ML approaches to commonly used methods. Finally, using UCE-derived SNPs and loci we validate species hypotheses using a standard delimitation method and a novel RF-based approach. We also demonstrate the utility of unsupervised ML on two previously published datasets.

## Materials and Methods

### Study System

For this study we utilized a short-range endemic (SRE; Harvey 2002) arachnid taxon in the order Opiliones (commonly called harvestmen). SRE taxa tend to have low dispersal ability and high ecological constraints, which leads to high population genetic structure and allopatric distributions, likely driven by niche conservatism (Wiens and Graham 2005). These biological characteristics make SRE taxa ideal candidates for species delimitation analyses, with high probability for new species discovery. Studies in SRE harvestmen (and SRE taxa in general) tend to show a great deal of underestimated diversity with numerous harvestmen species still being described even from well-studied areas (e.g., Derkarabetian and Hedin 2014; DiDomenico and Hedin 2016; Starrett et al. 2016; Emata & Hedin 2016).

The Pacific Northwest endemic genus *Metanonychus* Briggs, 1971 is a cryophilic harvestman that prefers moist forests, typically found underneath rotting logs/bark and in leaf litter. The genus and all species/subspecies were described by Briggs (1971), and currently includes three species: *M. idahoensis, M. setulus* with five subspecies (*setulus, mazamus, cascadus, navarrus*, and *obrieni*), and *M. oregonus* with two subspecies (*oregonus* and *nigricans*) (Fig. 1). *Metanonychus* is an ancient lineage; in a recent phylogenomic analysis of the superfamily Travunioidea (which contains *Metanonychus*), more genetic divergence is seen between the two samples of *Metanonychus* than in divergences between the vast majority of pairs of sister genera across all Travunioidea (Derkarabetian et al. 2018). Despite the ancient origin of this group, relatively few species were described, even though all “subspecies” are easily differentiated based on apparently fixed differences in male genitalic morphology (Briggs 1971). Recent systematic studies on related taxa corroborate the conservative nature of subspecies in these SRE harvestmen (Derkarabetian and Hedin 2014). As such, and more importantly, we consider *Metanonychus* species limits relatively straightforward where the species are “obvious” making this an excellent system to test ML approaches.

**Figure 1.**
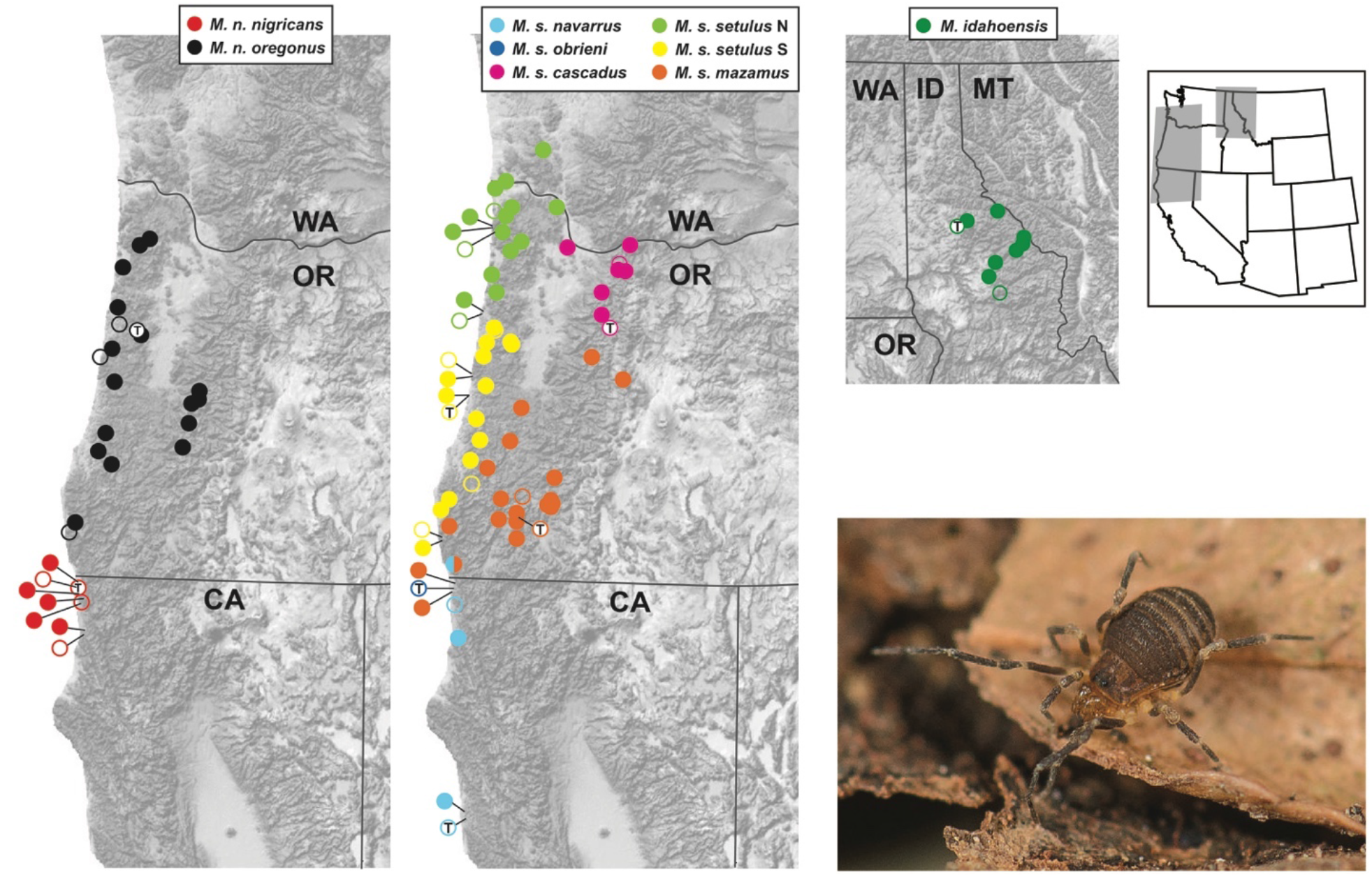
Geographic distribution of *Metanonychus*. Filled circles are collecting localities sampled for this study. Open circles are published records from Briggs (1971). Open circles with “T” indicate type localities. Live photo: *Metanonychus s. navarrus*.

### Species Delimitation Workflow

We consider species as “separately evolving metapopulation lineages” (de Quieroz 2007), that in practice are genetic clusters of samples corresponding to monophyletic lineages that show fixed morphological differences. For the discovery phase, our a priori species are based on inferred well supported COI clades and fixed differences in male genitalic morphology. We use two popular discovery-based genetic clustering approaches as “standards” to assess the utility and results of three ML methods. All SNP-based clustering analyses utilized the adegenet/STRUCTURE formatted file (.str) as input, which allowed minimal file format conversion from “standard” to ML approaches.

### Species Discovery

The COI gene was sequenced for at least one sample from every collecting locality, plus two outgroups from the sister genus *Sclerobunus*, using multiple primer combinations (online Appendix 1). DNA was extracted using the Qiagen DNeasy kit (Qiagen, Valencia, CA) using 2–3 legs, PCR experiments followed Derkarabetian and Hedin (2014) and amplified fragments were Sanger sequenced at Macrogen USA. The Sanger-sequenced COI dataset was supplemented with COI sequences derived as “UCE-bycatch” (e.g., Zarza et al. 2017; Hedin et al. 2018) for all UCE samples (see below). COI sequences were manually aligned and a phylogeny was reconstructed using RAxML v.8 (Stamatakis 2014) with 500 bootstrap replicates and the GTRGAMMA model. COI divergence dating was conducted with BEAST 2.4.8 (Bouckaert et al. 2014) using two calibrations: a strict clock calibrated at 0.0178 (Papadopolou et al. 2010), and a date calibration for the outgroups *S. nondimorphicus* (from coastal Oregon/Washington) and *S. idahoensis* (from Idaho), a well-known biogeographic break typically dated to 2–5 MY (Brunsfeld et al. 2001, and references therein), which was set to a uniform distribution of (2, 5).

The male genitalia in harvestmen tend to be species-specific and have been used in systematic studies across all taxonomic levels since the mid-1900s. We examined male genitalia for multiple samples of all described species/subspecies using standard scanning electron microscopy techniques. Images were taken using the FEI Quanta 450 FEG environmental SEM at the San Diego State University Electron Microscope Facility.

### Sequence Capture and SNPs

The number of studies utilizing UCEs in species delimitation and SNP-based population level analyses are increasing (e.g., Smith et al. 2013; Blaimer et al. 2016; Harvey et al. 2016; McCormack et al. 2016; Newman and Austin 2016; Zarza et al. 2016; Starrett et al. 2017; Hedin et al. 2018). Extractions were conducted as above, except in most cases whole bodies were used in digestions. Sequence capture of UCE loci followed the protocols available from the ultraconserved.org website and as in Starrett et al. (2017) and Derkarabetian et al. (2018) using the Arachnida 1.1Kv1 myBaits kit (Arbor Biosciences) designed by Faircloth (2017). Sequencing was done at the Brigham Young University DNA Sequencing Center on a HiSeq 2500 with 125 bp paired-end reads.

Raw reads were processed using phyluce (Faircloth 2005), adapter removal and quality control was done with an illumiprocessor wrapper (Faircloth 2013), and contigs were assembled with Trinity version r2013-02-25 (Grabherr et al. 2011). When matching contigs to probes, conservative values of 82 and 80 were used for minimum coverage and minimum identity, respectively, to filter potential non-target contamination (Bossert and Danforth 2018). Loci were aligned using MAFFT (Katoh and Standley 2013) and trimmed using gblocks (Castresana 2000; Talavera and Castresana 2007) with settings –-b1 0.5 –-b2 0.5 –-b3 10 –-b4 4. All loci were manually inspected in Geneious (Kearse et al. 2012) to fix obvious alignment errors and filtered for obvious non-homologs. Contigs corresponding to COI were identified by a local BLAST search in Geneious against available *Metanonychus* COI sequences. Although not used in species delimitation, a concatenated matrix of UCE loci with 70% taxon coverage was used to reconstruct a phylogeny using RAxML with 500 bootstraps and the GTRGAMMA model.

SNP datasets were created from sequence capture reads using published approaches (e.g., Zarza et al. 2017). The sample with the highest number of recovered UCE loci was used as a reference genome (*M. idahoensis*, OP2432). After adapter removal and quality control, reads for all samples were aligned to the reference using bwa (Li and Durbin 2009), the resulting SAM files were sorted using samtools (Li et al. 2009), PCR duplicates were identified and removed using picard (http://broadinstitute.github.io/picard), and all BAM files were merged. The Genome Analysis Toolkit 3.2 (GATK; McKenna et al. 2010) was used to realign reads and remove indels and SNPs were then recalibrated using “best practices” (van der Auwera et al. 2013). After recalibration SNPs were called and vcftools (Danecek et al. 2011) was used to create SNP datasets which varied in the percent of taxon coverage needed to include a SNP (50% and 70%). One random SNP from each locus was selected and the script adegenet_from_vcf.py (github.com/mgharvey/seqcap_pop) was used to create STRUCTURE-formatted (.str) files.

### Standard Genetic Clustering

As a comparison for the efficacy of unsupervised ML methods in inferring structure and optimal clustering, we used two popular approaches. First, STRUCTURE version 2.3.4 (Pritchard et al. 2000) was run for 1 million generations and 100,000 burnin on K values ranging from 2–10, with five replicates each. Structure Harvester (Earl and vonHoldt 2012) was used to determine optimal K via calculation of DK (Evanno et al. 2005) and Clumpak (Kopelman et al. 2015) was used to visualize output (http://clumpak.tau.ac.il/). Second, we used the adegenet R package (Jombart 2008; Jombart and Ahmed 2011) to conduct principal components analysis (PCA; dudi.pca function) and determine the optimal number of clusters and cluster assignment with discriminant analysis of principal components (DAPC) on scaled data.

### Unsupervised ML Visualization

Three unsupervised ML approaches were used for clustering (see Table 1). We executed RF through the randomForest R package (Liaw and Wiener 2002), extracting the scaled data from DAPC to a separate matrix. There are two important parameters associated with RF. The ntree parameter, the number of classification trees to create, was set to 5000. The mtry, the number of splits in the classification tree, was left at default for a classification analysis, which is square root the number of variables. The resulting proximity matrix was then used in both classic MDS (cMDS) and isotonic MDS (isoMDS). cMDS was executed using the MDSplot function in the randomForest package and isotonic MDS was conducted using the isoMDS function in the MASS R package (Venables and Ripley 2002).

VAE was implemented with a custom script utilizing the Keras python deep learning library (https://keras.io; Chollet 2015) and the TensorFlow machine learning framework (www.tensorflow.org; Abadi et al. 2015). As input for VAE we use SNP matrices converted via “one-hot encoding” where each nucleotide is transformed into four binary variables unique to each nucleotide (e.g., A = 1,0,0,0; C = 0,1,0,0; etc.) including ambiguities (e.g., M = 0.5,0.5,0,0) using a custom script. The VAE is composed of an encoder and a decoder. The encoder takes the one-hot encoded SNP data and infers the distribution of latent variables, given as a normal distribution with a mean (µ) and standard deviation (σ). The decoder then maps the latent distribution to a reconstruction of the one-hot encoded SNP data. As there are two latent variables, SNP data for each sample can be visualized as a reduced two-dimensional representation. Details of the VAE and the training procedure are in Supplementary File: Figure 1.

t-SNE was executed using the R package tsne (Donaldson 2016). After preliminary testing, several parameters were specified: maximum iterations (max_iter=5000), perplexity=5, initial dimensions (initial_dims=5), and number of dimensions for the resulting embedding (k=2). The maximum iterations value is relatively straightforward to determine as the KL divergence (a measure of the difference between high and low dimensional representations) should stabilize at a minimum. Perplexity is a measure of the balance between the local and global elements of the data; essentially how many neighbors a particular sample can have. This is a somewhat subjective parameter, where lower perplexity will produce tight well separated clusters, and higher values will produce more diffuse less distinguishable clusters. However, results and clusters are typically robust across a wide range of perplexity values (Pedregosa et al. 2011) and methods have been introduced to make perplexity selection automatic (Cao and Wang 2017). With large datasets it is recommended to perform dimensionality reduction on the data via PCA or a similar algorithm prior to implementing t-SNE (Pedregosa et al. 2011). As such, we perform t-SNE using the results of the initial PCA as input.

With RF and t-SNE, we also tested three different types of input format using the 70% SNP dataset. First, the SNPs were represented as raw nucleotides with ambiguities in standard IUPAC coding, extracted directly from .vcf files using the vcf2phylip script (github.com/edgardomortiz/vcf2phylip). Second, the raw SNPs were converted to haplotypes using the script SNPtoAFSready.py (github.com/jordansatler/SNPtoAFS). Third, the raw unphased nucleotides were converted into numerical format via one-hot encoding. For the first two datasets, the Ns were coded as blank, and PCA could not be conducted as the variables are categorical. As such, t-SNE was run using the cMDS output.

### Unsupervised ML Clustering

To assess the performance of clustering based on ML results relative to widely used STRUCTURE and DAPC approaches, four sets of clustering analyses were conducted using RF, VAE, and t-SNE outputs. First, to confirm that cluster assignments are equivalent to DAPC and STRUCTURE assignments, PAM clustering was conducted using the cluster R package (Maechler et al. 2018) with the optimal K selected from DAPC. The next three clustering methods test whether the optimal K can be inferred correctly relying solely on unsupervised ML results. PAM clustering was done on all output, including both the proximity matrix and cMDS for RF, across K of 2–10 with the optimal K having the highest average silhouette width (Rousseeuw 1987). Next, PAM clustering was conducted with the optimal K determined via the gap statistic using k-means clustering implemented in the factoextra R package (Kassambara and Mundt 2017). Finally, optimal K and clusters were determined via hierarchical clustering with the mclust R package (Scrucca et al. 2017) using only components retained via the broken stick algorithm implemented in the PCDimension R package (Coombes and Wang 2018).

### Species validation

We implement the commonly used Bayes Factor delimitation approach (*BFD; Leaché et al. 2014) with SNAPP (Bryant et al. 2012) using a 70% UCE SNP matrix created by the phyluce script “phyluce_snp_convert_vcf_to_snapp”. Multiple species hypotheses were tested based on current taxonomy, a priori species, ML clustering results, and an analysis where each individual specimen was treated as a unique species. SNAPP analyses were run with default settings for 100,000 generations, 10,000 burnin, and 48 steps. Each analysis was run twice to ensure consistency. Bayes Factors (Kass and Raftery 1995) were calculated (2 * log likelihood difference) to determine relative support of species hypotheses.

Next, we use the RF-based program CLADES (Pei et al. 2018), which uses Support Vector Machines, a type of supervised ML, to build a classifier based on labeled samples where samples are classified as either the same or different species. Several population genetic statistics are calculated for the simulated training data and the observed data, which are then treated as variables. The classifier is then used to infer whether the observed a priori species are equivalent to the same or different species. As input we use the UCE loci in two different analyses: 1) an analysis validating a priori species hypotheses (“spp” dataset); and 2) an analysis in which every individual was treated as a distinct species (“ind” dataset).

### Published Datasets

Uma notata *complex*. – Gottscho et al. (2017) explored lineage diversification and species limits in fringe-toed lizards of the *Uma notata* species complex, a group with a complicated taxonomic history. Using ddRAD data they find significant levels of gene flow between multiple species and determine that *U. rufopunctata* is a hybrid population. Several genetic clustering algorithms were used with differing results: DAPC favored an optimal K=5 (grouping the hybrid *U. rufopunctata* with *U. cowlesi*), while a model with admixture favored an optimal K=6 (splitting *U. scoparia* and showing varying levels of admixture for *U. rufopunctata* samples between *U. cowlesi* and *U. notata*). We reanalyzed their data with the intention of assessing unsupervised ML clustering/visualization in the face of significant gene flow and known hybrids. The published dataset with 597 SNPs was downloaded from Dryad (https://doi.org/10.5061/dryad.8br5c).

Phrynosoma coronatum *complex*. – The coast horned lizards of the genus *Phrynosoma coronatum* complex have received much attention with many species hypotheses put forth (summarized in Leaché et al. 2018). In an integrative approach Leaché et al. (2009) recover five well supported mtDNA clades that show little concordance with nuclear loci, ultimately integrating ecology and morphology to support three species (*P. blainvillii, P. cerroense*, and *P. coronatum*). More recently, Leaché et al. (2018) use SNP data coupled with *BFD testing all hypotheses derived from previous research, ranging from one to six species. A five species model is given the highest support, reflecting mtDNA and splitting *P. blainvillii* into three groups. Here, we use unsupervised ML methods for clustering, but more importantly to demonstrate their utility as a data visualization tool in a dataset showing high uncertainty in cluster probability assignments (fig. 1 of Leaché et al. 2018). Data were downloaded from dryad (https://doi.org/10.5061/dryad.k7k4m), and the SNP dataset in the .xml file was manually extracted and converted to .csv format for import into R.

## Results

### Species Discovery

*Metanonychus* specimens were collected from 79 different collecting localities. A total of 117 sequences were included in COI analyses (alignment length of 1182 bp); all new COI sequences have been deposited to GenBank (XXXX -XXXX). Seventy-seven sequences were acquired via Sanger sequencing and 38 were sequenced as UCE bycatch, with five samples being sequenced by both approaches, for a total of 110 *Metanonychus* specimens (plus two outgroups). UCE bycatch sequences possessed no stop codons, and for those samples sequenced via Sanger and as UCE bycatch, sequences were identical. COI divergence dating supports the ancient origin of this genus dating to ∼25 Ma (Supplementary File: Fig. 2). The RAxML phylogeny recovers a deep split between the “*nigricans* group” containing both subspecies of *M. nigricans* and the “*setulus* group” containing *M. idahoensis* and *M. setulus* with all subspecies. Each currently named taxon is monophyletic with bootstrap support values of 100 (Supplementary File: Fig. 3), except the *setulus* subspecies is polyphyletic separated into geographically cohesive northern and southern clades, although support for relevant internal nodes are weak.

**Figure 2.**
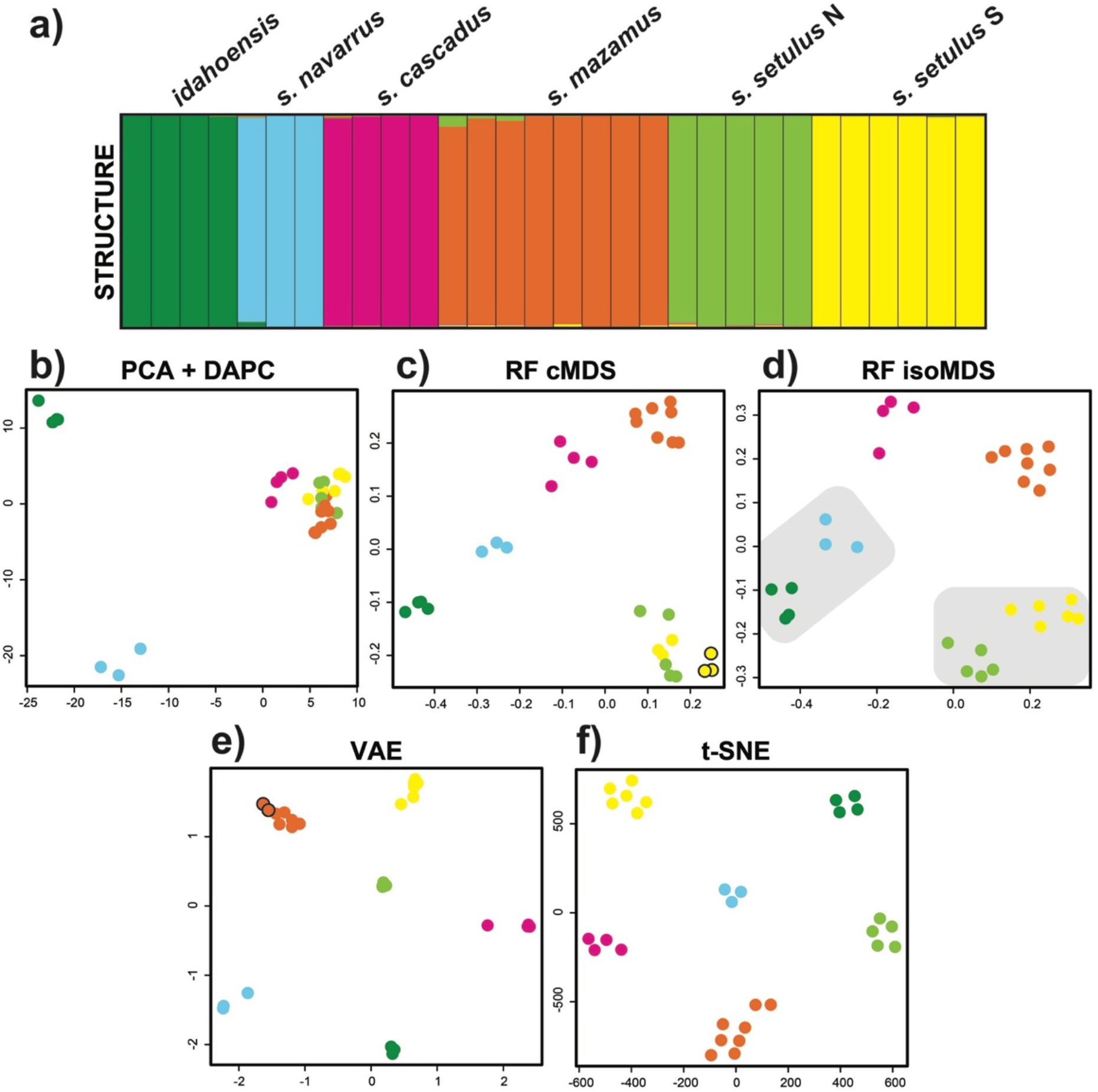
Clustering results for the *Metanonychus* 70% SNP dataset. a) STRUCTURE plot. b) PCA plot with DAPC clusters. c) random forest cMDS plot, all clustering algorithms favored K=6, except hierarchical clustering with K=7 (seventh cluster indicated with black outline). d) random forest isoMDS plot, all clustering algorithms favored K=6, except PAM clustering of RF output with K=4 (lumped clusters are indicated with grey shading). e) VAE plot, all clustering algorithms favored K=6, except hierarchical clustering with K=7 (seventh cluster indicated with black outline). f) t-SNE plot, all clustering algorithms favored K=6. cMDS = classic multidimensional scaling, isoMDS = isotonic multidimensional scaling.

**Figure 3.**
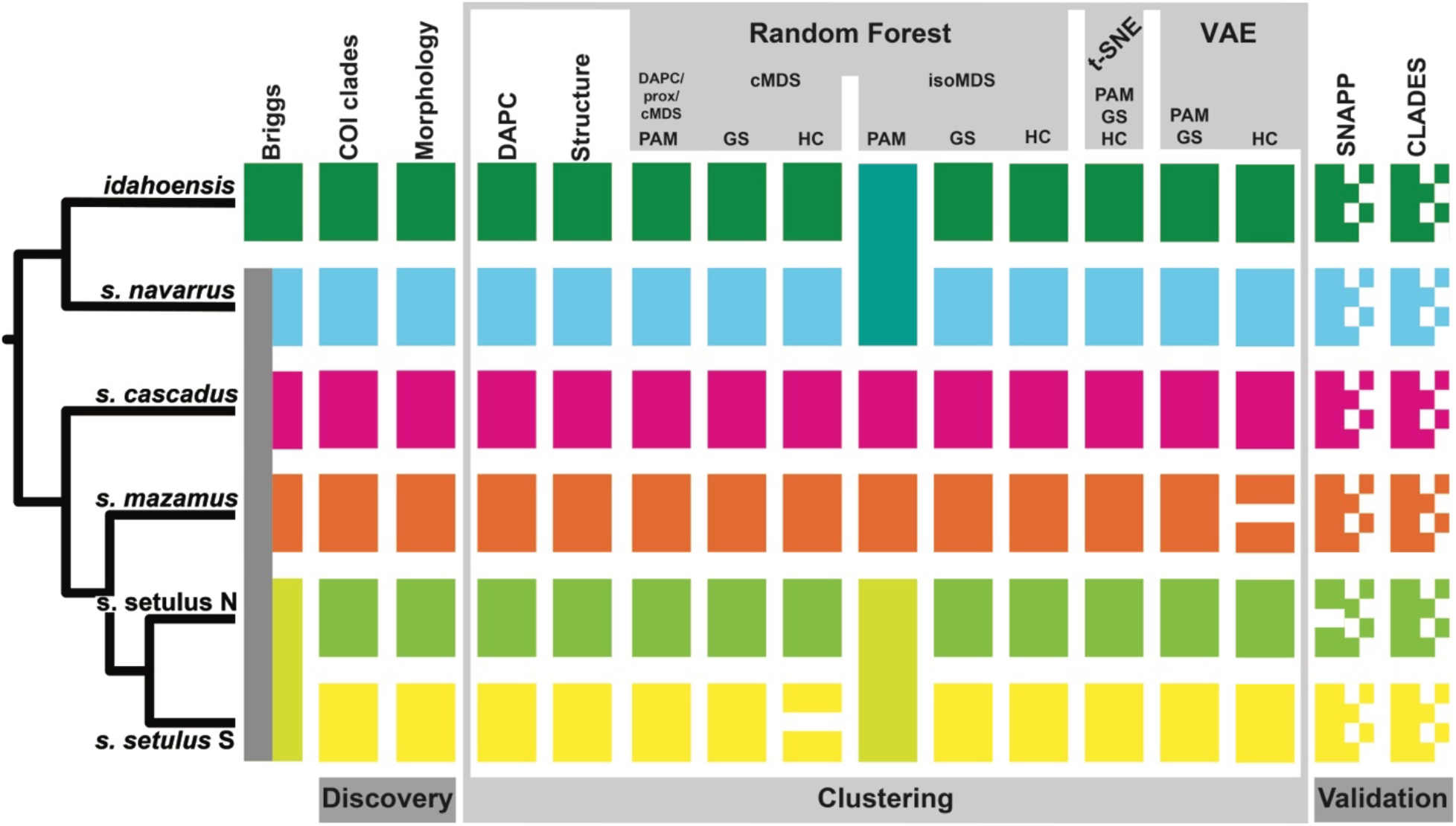
Integrative species delimitation results for the *Metanonychus* 70% SNP dataset. Species tree at left adapted from RAxML analysis of 70% UCE dataset. GS = gap statistic, HC = hierarchical clustering.

Male genitalic morphology show clear differences between all species/subspecies, including northern and southern clades of the *setulus* subspecies (Supplementary File: Fig. 4). Taken together, the discovery phase identified eight a priori species corresponding to the currently named species/subspecies (except *obrieni*, Appendix 1) with the *setulus* subspecies split into two genetically divergent, geographically cohesive clades with fixed differences in genitalic morphology.

**Figure 4.**
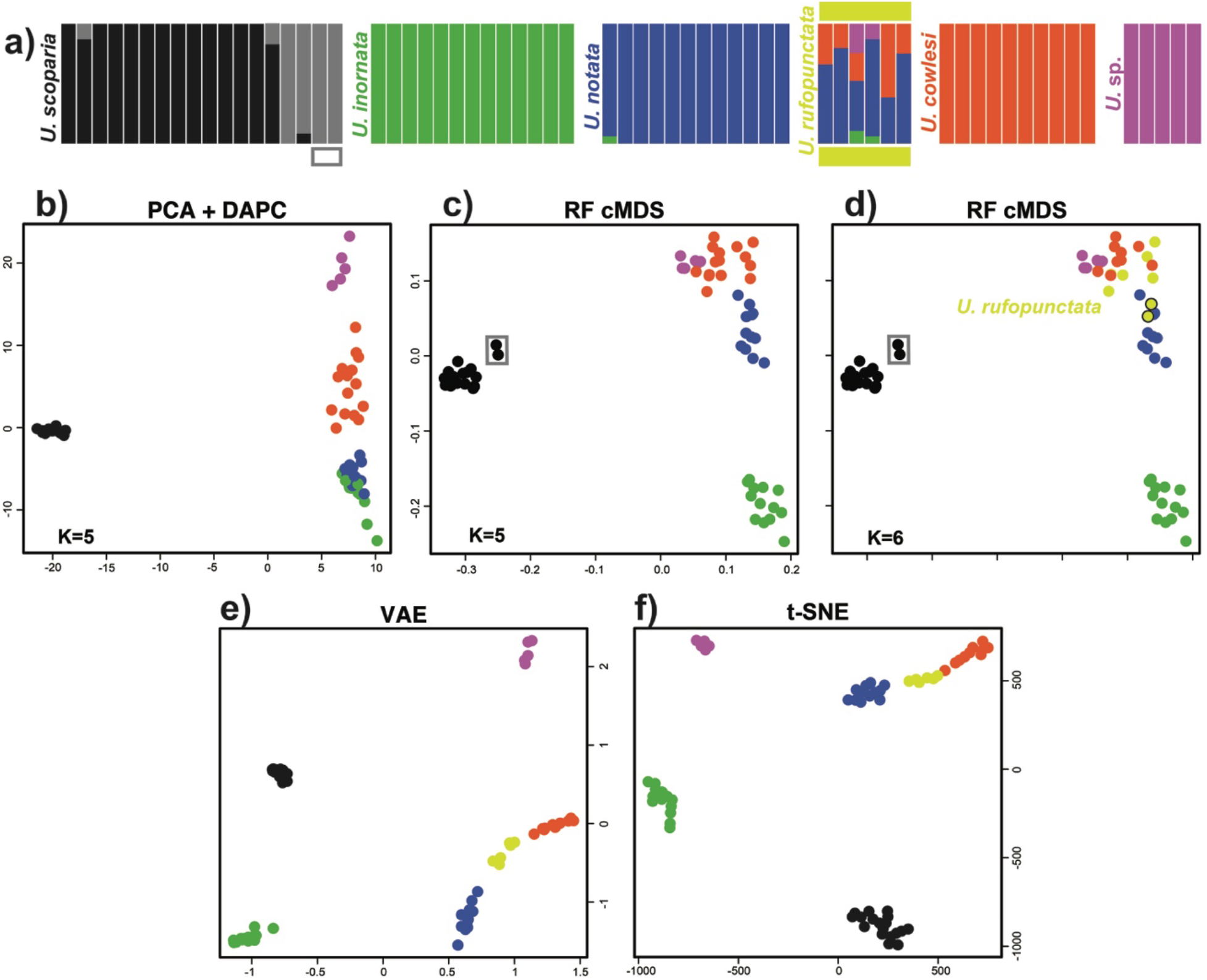
Clustering results for *Uma* dataset. a) STRUCTURE plot adapted from Gottscho et al. (2017). b) PCA with DAPC clusters. c) random forest cMDS plot with clusters identified via DAPC, PAM, and gap statistic. d) random forest cMDS plot with clusters identified via hierarchical clustering. e) VAE plot with K=6 a priori species. f) t-SNE plot with K=6 a priori species. Species are color coded as in Gottscho et al. (2017). Note: algorithmic clustering was only conducted on random forest output.

### Sequence Capture and SNPs

A total of 38 *Metanonychus* samples were included in UCE analyses, 36 of which were newly sequenced (online Appendix 1). Raw reads for sequence capture data have been deposited to SRA (XXXX). The 70% matrix included 185 loci (average of 158 per sample) with a mean locus length of 411 bp and a total length of 75,944 bp. The UCE phylogeny similarly confirms the monophyly of the *nigricans* and *setulus* groups and recovered the same clades as COI, but all internal nodes were fully supported (Supplementary File: Fig. 3). The *setulus* subspecies is recovered as monophyletic, albeit with reciprocally monophyletic northern and southern lineages. A 50% UCE matrix (278 loci, mean locus length of 384 bp, total length of 106,786 bp), produced an identical topology (not shown).

Due to the relatively high levels of divergence in *Metanonychus*, preliminary exploration of SNP datasets including all 38 samples resulted in datasets with too few loci or too sparse a matrix, with *M. nigricans* samples missing an average of ∼60% of SNPs (∼11% average samples in the *setulus* group). For the purposes of demonstrating ML clustering in *Metanonychus*, we focus on the monophyletic *setulus* group with six a priori species identified in the discovery phase. The *setulus* group included 30 samples and the 70% SNP dataset contained 316 SNPs (average of 250 per sample), while the 50% dataset contained 1263 SNPs (average of 774 per sample).

### Standard Genetic Clustering

For the 70% and 50% SNP datasets, both STRUCTURE (DK) and DAPC favored an optimal K=6 (Fig. 2 a, b; Fig. 3; Supplementary File: Fig. 5 and Fig. 6), recovering all six a priori *setulus* group species as distinct clusters, including the separate clades of the *setulus* subspecies.

### Unsupervised ML

Unsupervised ML analyses were relatively quick and computationally inexpensive taking 1–3 minutes for each of the three algorithms when run locally. All ML analyses were run multiple times producing identical clustering results. For the 70% dataset, all clustering approaches for RF (cMDS and isoMDS), VAE, and t-SNE resulted in an optimal of K=6, with the exception of the cMDS with hierarchical clustering resulting in an optimal of K=7 splitting the southern clade of the *setulus* subspecies, and hierarchical clustering of VAE with an optimal of K=7 splitting *mazamus* (Fig. 2, Fig. 3). Importantly, all K=6 clustering assignments were identical to those from DAPC and STRUCTURE. For the 50% dataset, an optimal of K=6 was found for the majority of analyses (Supplementary File: Fig. 5 and Fig. 6). However, the cMDS using hierarchical clustering resulted in K=7, splitting the northern clade of the *setulus* subspecies, and hierarchical clustering of VAE resulted in K=7, splitting *mazamus*. Clustering of the 50% dataset based on isoMDS was more variable, with an optimal K=4 for hierarchical clustering and K=1 for the gap statistic. All VAE and t-SNE clusters were obvious. VAE clusters were robust, being recovered identically across five replicate analyses, and clear separation between clusters is seen when σ (standard deviation) is included (Supplementary File: Fig. 7). t-SNE clusters were robust to perplexity values from 5–25, after which samples became randomly dispersed (Supplementary File: Fig. 8). The unsupervised ML approaches produced plots with easier interpretability relative to PCA, with species clusters showing more separation in two-dimensional space. Similar plots for RF and t-SNE were obtained using input where SNPs were coded in multiple ways (Supplementary File: Fig. 9).

### Species Validation

*BFD showed increasing likelihood with increasing species (Table 2), with Bayes Factors heavily favoring the analysis in which all individual specimens were treated as distinct species (K=30). Only considering hypotheses recovered in the discovery phase, the “7N” species hypothesis was favored, recognizing all six a priori species plus two species in the northern clade of the *setulus* subspecies. CLADES requires that each locus have data for at least one sample within every a priori species. As a result, the “spp” dataset had 177 loci and the “ind” dataset had 12 loci. CLADES supported the species status of all six a priori species. However, species status was also supported when each sample was treated as a distinct species (Fig. 3).

**Table 2.** Results of *BFD hypothesis testing.

| Species | Justification | A | B | Bayes Factor |
| --- | --- | --- | --- | --- |
| 2 | Briggs' species | -3674.33 | -3885.74 | ~6924 |
| 4 | 70% isoMDS PAM, 50% isoMDS HC | -2917.94 | -2910.14 | ~5192 |
| 5 | Briggs' species + subspecies | -2384.94 | -2386.83 | ~4135 |
| 6 | a priori species | -2210.48 | -2211.17 | ~3785 |
| 7 M | split *s. mazamus*: VAE HC | -2135.1 | -2136.23 | ~3635 |
| 7 N | split *s. setulus* N: 50% cMDS HC | -1797.95 | -1798.71 | ~2960 |
| 7 S | split *s. setulus* S: 70% cMDS HC | -2165.62 | -2166.25 | ~3695 |
| 30 | all individuals | -320.3 | -316.24 | - |

### Supplementary Material

All *Metanonychus* input matrices (COI, UCE SNPs, UCE loci, and .csv files) are available from the Dryad Digital Repository: http://dx.doi.org/10.5061/dryad.[NNNN]. Resulting phylogenies are available via TreeBASE (XXXX). Two custom scripts were created to run ML analyses: an R script to run random forest, t-SNE, and all clustering algorithms (github.com/shahanderkarabetian/uml_species_delim), and a python script to run VAE (github.com/sokrypton/sp_deli).

### Published Datasets

Uma notata *complex*. – All clustering based on RF with cMDS favored a K=5 scenario, with cluster assignment identical to DAPC results, with the exception of hierarchical clustering favoring an optimal K=6 (Fig. 4). In this case, a distinct cluster was identified for all *U. rufopunctata* and two samples of *U. notata*. The optimal of K=6 recovered in Gottscho et al. (2017) does not differentiate *U. rufopunctata*, instead splitting *U. scoparia*. The cMDS plots do show two somewhat distinct samples of *U. scoparia*, which correspond to samples placed in the sixth cluster. Clustering results ranged from K=4 in PAM, lumping *U. cowlesi, U. notata*, and *U. rufopunctata*, to K=7 in some replicates of t-SNE clustered with gap statistic splitting *U. scoparius*. The t-SNE and VAE plots recover the hybrid species *U. rufopunctata* as a linear “grade” between the parental species *U. cowlesi* and *U. notata*, and assignment uncertainty of the hybrid samples are seen when σ is also visualized (Supplementary File: Fig. 7).

Phrynosoma coronatum *complex*. – As expected, clustering via DAPC, RF, VAE, and t-SNE with nuclear SNP data did produce groups congruent with mitochondrial clades, with the exception of *P. coronatum*(Fig. 5). DAPC favored K=4 (*P. coronatum*, southern *P. cerroense*, northern CA *P. blainvillii*, and northern P*. cerroense* + the rest of *P. blainvillii*), while PAM clustering favored K=2. The differing cluster assignments of *P. cerroense* lineages reflects their polyphyly in the SNP phylogeny of Leaché et al. (2018). While all plots arrange samples in a way reflective of their genetic similarity, the more diffuse spatial arrangement of samples in the t-SNE embedding and the σ of the VAE are particularly informative and reflective of cluster probability assignments for *P. blainvillii* samples (Fig. 5d and Supplementary File: Fig. 7).

**Figure 5.**
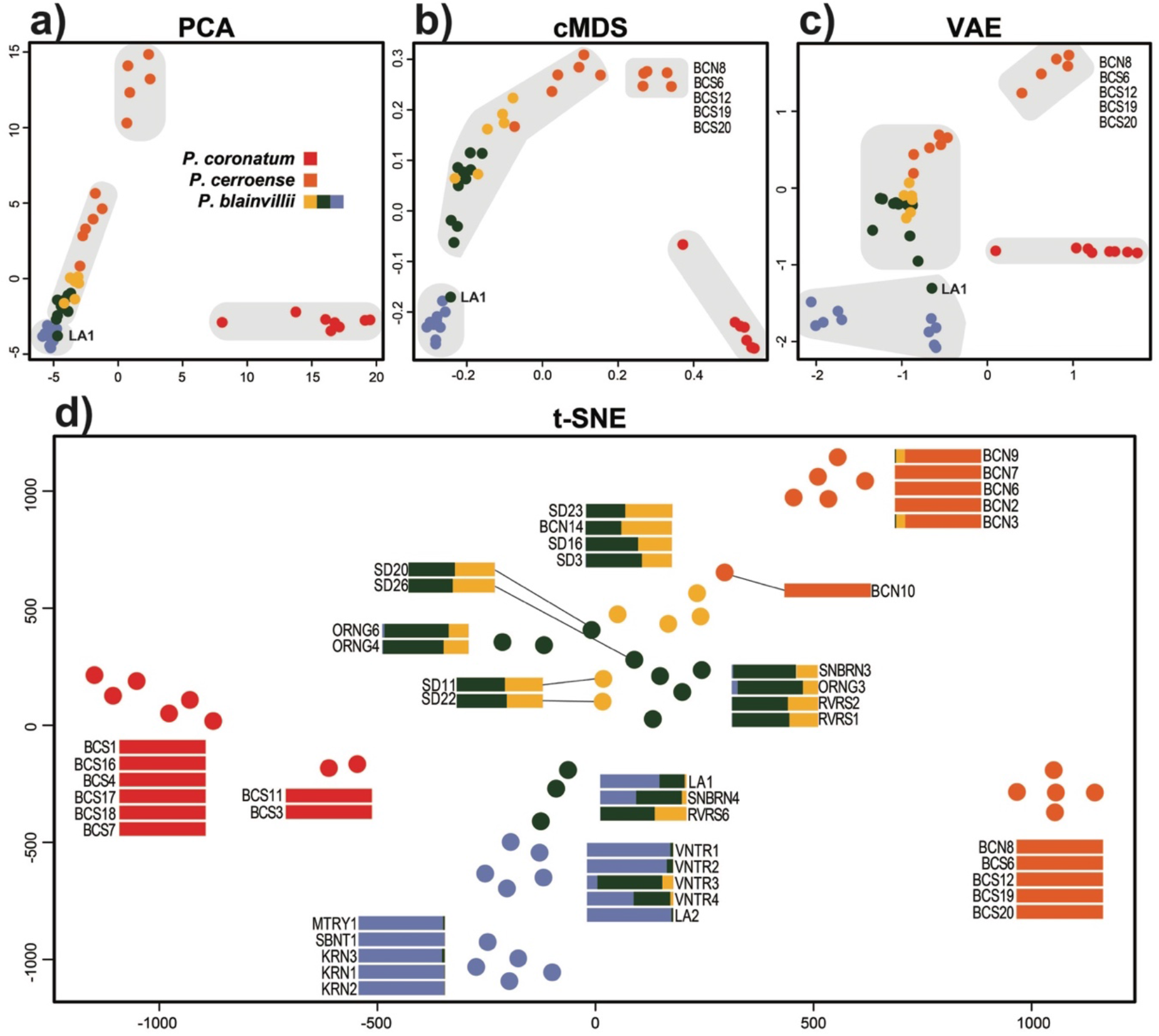
Clustering results for *Phrynosoma* dataset. a) PCA plot. b) random forest cMDS plot. c) VAE plot. For parts a-c) samples are colored by mtDNA clades recovered in Leaché et al. (2009), and grey boxes indicate optimal clustering of K=4 recovered via DAPC. d) t-SNE embedding, with corresponding assignment uncertainty for each sample adapted from Leaché et al. (2018). Samples are color coded as in Leaché et al. (2018).

## Discussion

### Reconsidering (SRE) Species Delimitation

Commonly used validation approaches relying on genomic-scale data have the potential to identify population structure and oversplit taxa (e.g., Sukumaran and Knowles 2017), a problem that can be exacerbated when studying SRE taxa with inherently high levels of population structure. Model-based validation analyses relying on the multispecies coalescent as currently implemented (e.g., BPP, SNAPP) seek to identify separate panmictic gene pools. This approach may not be suitable for *all* taxa given the diversity of biological characteristics unique to particular groups or organismal types with differing degrees of population structure and isolation, etc. (Sukumaran and Knowles 2017). While the issue of population structure in species delimitation has recently come under focus from a methodological perspective, the potential misinterpretation of population structure as species level divergences in empirical data has been a concern for taxonomists focusing on SRE taxa for a relatively long time (e.g., Hedin 1997), and continues to be so (e.g., Boyer et al. 2007; Bond and Stockman 2008; Niemiller et al. 2012; Barley et al. 2013; Satler et al. 2013; Fernández and Giribet 2014; Hedin 2015; Hedin et al. 2015).

Unsupervised ML clustering of SNP data provided reasonable species hypotheses that were largely identical to commonly used discovery-based analyses. However, when used with validation methods, the same data supported unrealistic results severely overestimating the number of species. Most importantly, clusters identified in unsupervised ML approaches obviously correspond to species, implying that cluster separation was dominated by species-level divergences and not population structure. If validation analyses show increasing support for more complex species delimitation models, up to the most unrealistically complex model possible given the data (i.e., each individual specimen as a distinct species), those analyses do not contribute useful information to the final species hypotheses. Similarly, the possibility of the most complex model being favored, whether actually tested or not, makes “support” for any less complex alternative models meaningless. If we did not run the K=30 SNAPP analysis or the “ind” analysis in CLADES, a more realistic 6–7 species hypothesis would be favored validating all a priori species, without any consideration of more complex hypotheses that are actually more likely. For *Metanonychus*, validation analyses were effectively ignored in the formation of final species hypotheses, and the information content of the SNP dataset was squandered, not being used to its full potential. While *BFD/SNAPP is useful for testing alternative assignment hypotheses, its use as a validation tool to determine the number of species is certainly problematic for SRE taxa, and more broadly for any taxon with significant population structure.

Because model-based validation analyses have the potential to delimit population level divergences, that does not mean they *only* identify population-level divergences. However, the confirmation that validation analyses are operating at the species level can only be assessed when species delimitation is conducted in an integrative framework, and we reiterate the statement by Sukumaran and Knowles (2017) that external information (i.e., different data types) are needed to confirm delimitations made based on genetic-only analyses. Ultimately, we argue that the separation of empirical species delimitation into two distinct phases (discovery and validation) limits the potential utility of the “validation” data type in informing species hypotheses in a truly integrative manner. Data types used in the discovery phase inform the a priori species hypotheses used as input for the validation phase, but the data type used in validation does not reciprocally inform the other data types. Ideal integrative taxonomy as described by Schlick-Steiner et al. (2010) utilizes multiple data types in a reciprocally illuminating framework where discordance between datasets requires consideration of the underlying biological processes.

### Machine Learning in Species Delimitation

The goal of this study was to explore how well unsupervised ML methods can successfully identify clusters equivalent to species and correctly infer the expected number of clusters. We argue that species delimitation in *Metanonychus* was relatively “simple” showing essentially no discordance between datasets and provided an excellent study system to explore novel approaches. In an integrative framework, our results suggest that the expected number of species, determined via mitochondrial and morphological analyses, can be correctly inferred across multiple clustering algorithms using the RF distances, the latent variables of VAE, and the t-SNE embeddings. Most importantly, unsupervised ML approaches coupled with standard clustering algorithms did not oversplit the data by distinguishing samples based on population-level structure, but instead formed clear clusters equivalent to species-level divergences. While these unsupervised approaches seemingly work well with relatively clear species, their ability to correctly cluster samples in more difficult speciation scenarios (e.g., rapid and recent divergence, divergence with gene flow, etc.) remains to be tested, although results in *Uma* are promising.

For unsupervised RF, more consistent and “accurate” clustering was achieved using the cMDS output. Like DAPC, multiple dimensions are used to inform the optimal clustering strategy. Conversely, isoMDS by default only outputs two dimensions for clustering. isoMDS may be suitable for significantly diverged taxa, in which case it can sometimes produce a better two-dimensional visualization of the data relative to cMDS. VAE and t-SNE clusters were exceedingly obvious regardless of data type, and robust across multiple iterations and varying parameters. t-SNE was designed purely for the visualization of high dimensional data, although given a low dimensional embedding as output, clustering is an obvious application. It has been noted that t-SNE clusters, cluster size, and distances between clusters may not have any relevant meaning (Wattenberg et al. 2016) and clusters should be interpreted with caution. As t-SNE does not preserve the density of actual clusters completely, density-based clustering algorithms (Ester et al. 1996; Campello et al. 2013) may offer an improvement relative to other clustering approaches. Regardless, in the datasets used here, inferred clusters have obvious biological meaning corresponding to species which were corroborated by other analyses and data types. More consistent and accurate clustering results were obtained with the 70% taxon coverage dataset. Samples with a higher percentage of missing data might be reconstructed in closer proximity by unsupervised ML methods, regardless of phylogenetic proximity, simply because they share high levels of missing data. This is particularly the case with data converted to one-hot format where a missing SNP was coded as “0,0,0,0”, although we designed our VAE to mask missing data.

Neural networks have mostly been designed/used for identifying the latent space of images, the most relevant examples including the citizen science natural history observational platform iNaturalist (www.inaturalist.org) and classification of ants (Boer and Vos 2018). Here we show that VAEs, which leverage neural networks to learn a probability distribution of the data, can learn phylogenetic structure with the latent variables. In contrast to t-SNE, VAEs are nicely derived from formal Bayesian probability theory, and can hence be used to score the probability that the new data belongs to a trained set of data or is a new species. The standard deviation around samples/clusters is an inherent result of a VAE analysis and visualization makes the assessment of cluster distinctiveness or uncertainty relatively straightforward. One drawback is that it is not straightforward when to stop training a VAE. Overtraining a VAE can lead to overfitting the data, which results in clusters that are still present, but the probability distribution over the data is less general, and hence cannot be used reliably for downstream analysis. One solution is to partition a small fraction of the training data as a validation set, which can be used to determine when training should be stopped, a technique in ML known as early stopping (Goodfellow et al. 2016), although we use a “dropout” approach to prevent overfitting. Given results presented here, the robustness of output to parameter variation, and its Bayesian nature, VAEs are very promising for future incorporation into systematic applications.

Data visualization is an important aspect of empirical research. With genetic data, whether used as loci or SNPs, this can be in the form of a phylogeny or via a dimensionality reduction method. Regardless of whether downstream clustering is performed, unsupervised ML methods like t-SNE and VAE offer excellent options for relatively quick and informative data visualization that can help examine uncertainty in a priori groupings or recognize misidentifications and paraphyly, both of which are problematic for species hypotheses if data are destined for downstream model-based analyses. The placement of hybrid populations of *Uma* and the arrangement of assignment uncertainty in *Phrynosoma* are displayed in low-dimensional space in spatially meaningful ways. The recently developed Uniform Manifold Approximation and Projection method (McInnes and Healy 2018) is a dimensionality reduction technique similar to t-SNE but with numerous benefits including better preservation of global structure and potential embedding in larger dimensional space benefitting downstream clustering.

Unsupervised ML methods do not make assumptions about data type (e.g., genetic versus morphological, etc.); data are merely treated as data. If approaches that are not specifically designed for a particular data type successfully identify/corroborate a priori species, the resulting species decisions are more robust. However, the underlying assumption is that the analyses are operating at the species level. As with many dimensionality reduction techniques, unsupervised ML methods will uncover any underlying structure regardless of the taxonomic level or type of data. As such, integrative taxonomy with multiple data types and analytical approaches is ideal. Conversely, this insensitivity to taxonomic scale makes unsupervised ML relevant to population level analyses and phylogeography as well as species delimitation in taxa across varying divergence times, for example, divergences of ∼20 Ma in the *Metanonychus setulus* group down to much more recent species divergences of <1 Ma reported for *Uma*(Gottscho et al. 2017).

An additional appeal of some ML approaches is their ability to be conducted in a “semi-supervised” manner, where some samples can be labeled (e.g., assigned to a species) while others are left unassigned. For example, semi-supervised analyses could be used for species assignment of samples with unknown determination, like females of *Metanonychus*, or in taxa where the vast majority of specimens are known from juveniles that cannot be identified to species (e.g., Hedin et al. 2018). While fully supervised approaches have been used for this same reason, for example with COI barcoding (e.g., Weitschek et al. 2014; Archer et al. 2017), utilizing a semi-supervised ML approach (e.g., McInnes and Healy 2018) saves the need for creating a training dataset and associated assumptions. In either case, given the increasing incorporation of museum specimens in genomic analyses (McCormack et al. 2016; Blaimer et al. 2016; Ruane and Austin 2017; Sproul and Maddison 2017) it is now feasible to directly include type specimens in species delimitation. In the case of semi-supervised methods, type specimens (or specimens from type localities, etc.) can be included in analyses as labeled data while all other samples are left unlabeled, or in a supervised approach, data from type specimens could be used in training dataset construction.

If model testing is integral to the study it seems more logical, particularly in cases where genetic data is the only reliable way to assess species limits (i.e., cryptic species), to rely on algorithms that utilize prior information in the form of training data based on parameters associated with the particular biological characteristics of a given organismal type, thus taking the biology of the organism more directly into account. For potential future analyses of SRE harvestmen using supervised ML methods, training data could consist of multiple “curated” SRE datasets where species are known and well-supported, which would then be used for SRE taxa with unknown or uncertain numbers of species. While CLADES oversplit *Metanonychus* supporting every individual as a species, we do not see this as a negative for the approach, but rather as imperative to create and use curated training datasets reflecting the biological characteristics of the study organism to fully leverage the power of this approach. More recently, Smith and Carstens (2018) developed delimitR, a supervised ML approach that treats species delimitation as a classification problem, using the binned multidimensional Site Frequency Spectrum as the predictor variable to build an RF classifier that can distinguish among different speciation models, the response variables, selecting the model with the most votes. Training data is simulated based on specification of several priors (guide tree, population size, divergence time, migration) either known or estimated for the particular study system. DelimitR is a promising approach as priors are used to create the simulated data for classifier construction, making the analysis more specific to the biology of the focal taxon.

In general, unsupervised ML approaches offer the benefits of better data visualization in two-dimensional space and the ability to accommodate various data types. Like current methods combining multiple data types into a single analysis (e.g., Guillot et al. 2012; Solis-Lemus et al. 2015), it may be feasible to do an integrative unsupervised ML analysis where various data types (e.g., morphological, genetic, chemical profiles, etc.) are combined into a single dataset for downstream clustering. Many ML algorithms are well-suited for species delimitation, providing promising avenues of incorporation into standard systematics protocols and excellent resources are available for implementation (e.g., http://scikit-learn.org, https://keras.io, www.tensorflow.org). ML algorithms, even those designed for image analysis or pattern and text recognition, all seek to identify and learn the underlying structure of input data via dimensionality reduction of some form. This can be leveraged for all data types in diverse ways, for example, representing a multidimensional vector of population genetic statistics as an image to be analyzed via neural networks (Kern and Schrider 2018). As recently discussed in regard to population genetics (Schrider and Kern 2018), with a basic understanding of the types of ML algorithms, the applications to species delimitation become obvious and exciting with the potential to aid in all aspects of systematic biology.

### Learning from Metanonychus

Multiple data types and analytical approaches favor six species in the *setulus* group, providing robust final species hypotheses. Although some analyses favored more than six species, we prefer more conservative species hypotheses that are robust to data and analysis type (e.g., Carstens et al. 2013). As a result of integrative species delimitation, we elevate all subspecies of the *setulus* group to full species, now consisting of *M. idahoensis, M. navarrus* **new comb**., *M. cascadus* **new comb**., *M. mazamus* **new comb**., and *M. setulus*. In addition, all analyses supported the northern clade of the *setulus* subspecies as a distinct species, which we describe as ***M. xxxx* n. sp**. Derkarabetian and Hedin (Appendix 1). Based on examination of type specimens, *M. obrieni* is synonomized with *M. navarrus* (Appendix 1). The *nigricans* group had too few samples for reliable clustering when analyzed alone. However, both the morphological divergence seen in male genitalia and nuclear divergence supports elevating the *M. nigricans* subspecies to full species: *M. nigricans*, and *M. oregonus* **n. comb**. Based on our results, we reiterate that the subspecies rank common in several groups of SRE harvestmen are conservative estimates considering these “subspecies” also show fixed morphological differences that were used for the initial diagnosis.

*Metanonychus* is a relatively ancient genus, persisting in mesic forests of the Pacific Northwest since the late Oligocene, and its species are relatively old dating up to ∼10 Ma with extremely high levels of population divergence. From a biogeographical perspective, it is interesting to note that *M. idahoensis* from northern Idaho is recovered as sister to *M. navarrus* from northern California, to the exclusion of all taxa from Oregon and Washington. The break between mesic forests of Idaho and coastal Oregon/Washington is found in numerous taxa typically attributed to the formation of the Cascades dating to 2–5 Ma (Brunsfeld et al. 2001, and references therein). Divergence dating analyses here estimate that the split between *M. idahoensis* and *M. navarrus* is much older, dating to ∼12 Ma (average K2P-corrected COI divergence of 16.8%) suggesting the possibility of an older connection and divergence between these regions. Further exploration of this result in the context of Pacific Northwest biogeography is needed (e.g., Brunsfeld et al. 2001, Steele et al. 2005; Carstens and Richards 2007). These results reaffirm the importance of SRE taxa and their inclusion in exploring and elucidating, sometimes unexpected, patterns of regional biogeography and geologic history (e.g., Boyer and Giribet 2009; Hedin et al. 2013; Emata and Hedin 2016).

## Acknowledgements

For assistance during fieldwork we thank Casey Richart, James Starrett, Alan Cabrero, Erik Ciaccio, and Morganne Sigismonti. The initial inspiration for this work was provided by Ty Roach. We thank Nick Vinciguerra for help during SNP processing and Morganne Sigismonti for assisting with initial morphological examinations. Type specimen loans were kindly provided by Darrell Ubick (California Academy of Sciences). An earlier version of the manuscript was improved through discussion with Jeet Sukumaran.

